# Accurate inference of population history in the presence of background selection

**DOI:** 10.1101/2024.01.18.576291

**Authors:** Trevor Cousins, Daniel Tabin, Nick Patterson, David Reich, Arun Durvasula

## Abstract

All published methods for learning about demographic history make the simplifying assumption that the genome evolves neutrally, and do not seek to account for the effects of natural selection on patterns of variation. This is a major concern, as ample work has demonstrated the pervasive effects of natural selection and in particular background selection (BGS) on patterns of genetic variation in diverse species. Simulations and theoretical work have shown that methods to infer changes in effective population size over time (*N*_*e*_(*t*)) become increasingly inaccurate as the strength of linked selection increases. Here, we introduce an extension to the Pairwise Sequentially Markovian Coalescent (PSMC) algorithm, PSMC+, which explicitly co-models demographic history and natural selection. We benchmark our method using forward-in-time simulations with BGS and find that our approach improves the accuracy of effective population size inference. Leveraging a high resolution map of BGS in humans, we infer considerable changes in the magnitude of inferred effective population size relative to previous reports. Finally, we separately infer *N*_*e*_(*t*) on the X chromosome and on the autosomes in diverse great apes without making a correction for selection, and find that the inferred ratio fluctuates substantially through time in a way that differs across species, showing that uncorrected selection may be an important driver of signals of genetic difference on the X chromosome and autosomes.

## 2 Introduction

Understanding how effective population size has changed in the past – that is, reconstructing the population size trajectory *N*_*e*_(*t*) – is crucial in order to understand the evolutionary history of any species [1]. Coalescence-based approaches to inferring *N*_*e*_(*t*) are attractive due to their low sample size requirements (as few as two chromosomes) [2, 3, 4, 5]. These methods leverage the density of local heterozygosity to estimate the time since the most recent common ancestor (TMRCA) at each location along the genome, which is used to infer *N*_*e*_(*t*). However, these methods assume erroneously that loci across the genome evolve neutrally, despite the evidence of profound effects of linked natural selection on patterns of genetic variation in many species, including humans [6, 7, 8, 9, 10, 11, 12, 13, 14, 15, 16].

Simulation and theoretical studies have provided compelling evidence that selection biases inferences of *N*_*e*_(*t*) [17, 18, 19, 20]. Using simulations, Schrider et al. [18] demonstrated that in the presence of a selective sweep, the PSMC underestimates true *N*_*e*_(*t*) in times more recent than the onset of selection. This effect becomes greater as the frequency or intensity of sweeps increases. Johri et al. [19] simulated a model of widespread linked selection, and showed increasingly inaccurate estimates of *N*_*e*_(*t*) at all time scales whether PSMC [2], MSMC [3], or fastsimcoal [21] was used for inferences. Using an analytical approach that models the effects of linked selection as a rescaling of *N*_*e*_(*t*) by a locus-specific constant, Boitard et al. [20] confirmed a spurious decrease in inferred *N*_*e*_(*t*) that comes from not accounting for these effects *N*_*e*_(*t*). However, the impact of BGS on inferring *N*_*e*_(*t*) on real data remains unknown, and it is unclear how to obtain accurate estimates of this quantity in the presence of BGS.

Hudson and Kaplan [22] and Nordborg et al. [23] introduced a model to approximate the effects of BGS, by scaling local genomic *N*_*e*_ to account for loss of diversity due to linkage with deleterious alleles. Motivated by this approximation, McVicker et al. [6] explicitly estimated *b*_*i*_, the fraction of expected neutral diversity at site *i*, across the human genome. They estimate that linked selection results in a genome-wide average reduction in diversity of 19–26% on the autosomes, and 12-40% on the X chromosome. Primarily, they attributed this reduction to background selection (BGS) – the loss of genetic diversity due to linkage with alleles under purifying selection – but could not rule out a contribution from selective sweeps. Later work provided evidence that selective sweeps had little effect on diversity levels in humans [7], supporting the interpretation of these patterns as largely driven by BGS - though this has been contested [24]. By leveraging whole-genome sequencing data from the 1000 Genomes project [25] and more detailed functional annotations, Murphy et al. [26] re-estimated the contribution of linked selection in shaping human genetic diversity. They generated a much-improved B-map relative to the earlier map of McVicker et al. [6], producing a set of genome wide *b*_*i*_ values describing the strength of BGS. They showed that this map is sufficient to explain ∼60% of the variance in autosomal diversity levels at the megabase scale. They also concluded that selective sweeps have little or no effect on linked neutral diversity.

The demonstrated large effect of BGS on inferring *N*_*e*_(*t*) [18, 19, 20], combined with the empirically large demonstrated impact of BGS in diverse species including humans [6, 26], raises questions about whether inference of population size changes from genetic variation data under the assumption of neutrality is reliable. Here, we extend the PSMC algorithm to handle local changes in the mutation or expected coalescent rate in a software package we call PSMC+. We adopt a first-order approximation to the effects of BGS as a rescaling of population size with a locus-specific constant [22, 23]. Using forward-in-time simulations which explicitly incorporate selection, we demonstrate our approach is largely accurate even if BGS is widespread. We also test applications of the method when there is incomplete information about the strength of background selection across the genome, as is the case for most species. We test the accuracy of coalescent time inference in the presence of background selection and find that PSMC+ is accurate even in its presence. PSMC+ can also be used to obtain unbiased inferences even in the presence of variation in the mutation rate over the genome.

## 3 New Approaches

PSMC [2] models the density of heterozygous positions between two haploid genomes or within a single diploid genome, which reflects mutations that have accumulated since the two haploid genomes shared a common ancestor, to infer the time since the most recent common ancestor (TMRCA). Then, it uses this distribution to infer a piecewise constant effective population size trajectory through time, *N*_*e*_(*t*), integrating information over all loci in the genome. To model the probability of observing a heterozygous position, PSMC uses the population mutation rate *θ* = 4*N*_*e*_*µ* – where *N*_*e*_ is the local genomic effective population size, and *µ* is the mutation rate per base pair per generation, which PSMC assumes is constant across the genome. However, *µ* is known to vary across the genome [27, 28], and the effects of BGS are frequently modeled as variations in *N*_*e*_ [23, 22]. *Both of these effects violate PSMC’s assumption of constant θ*, raising the potential of bias in inference of *N*_*e*_(*t*), which we confirm in this work.

Here, we explore three approaches to overcoming bias in inferring *N*_*e*_(*t*) in the presence of background selection. First, we use a high resolution map of BGS to adjust the emissions model of PSMC. Second, we use a low resolution map of BGS based on distance to exon to adjust the emissions model of PSMC. Third, we use post-hoc scaling to adjust the output of PSMC based on the heterozygosity at the top 1% of sites furthest from exons. The last two approaches are useful for species where high resolution maps of BGS do not exist.

In detail, at a locus *i*, PSMC models *m*_*i*_, the number of segregating sites at locus *i*, which given a coalescent time, *t*_*i*_, can be written as

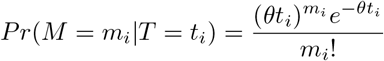

We modified the PSMC framework to condition on local variations in *θ* using a factor *f*_*i*_, which eliminates the bias in inferring *N*_*e*_(*t*) if the true factor is known

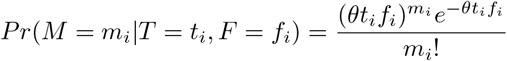

Further details are provided in the Methods section. We refer to our modified framework as PSMC+ and have released open-source software implementing the method (see Code Availability).

## 4 Results

### 4.1 Simulations

We performed forward-in-time simulations using SLiM v3 [29] with a realistic exon map, distribution of fitness effects, and recombination rates (see Methods) (Supplementary Figure 1). To correct for BGS in these simulations, we used a B-map based on genetic distance to simulated exons, which while not as optimized as the one developed by Murphy and colleagues for real data, allowed us to explore the behavior of PSMC+ (see Methods). We simulated both a constant population size demographic and a demographic history of population size changes similar to that as inferred in humans. We then compared *N*_*e*_(*t*) as inferred from PSMC+ or PSMC. In what follows, we only use PSMC+ where we are inputting into the model a map of heterogeneous rates; we use “PSMC” to refer to our implementation of the original algorithm that assumes homogeneous rates across the genome.

We find that PSMC+ is unbiased for simulations of a constant effective population size (Figure 1a), as expected based on the approximation of [22, 23] that describe the effects of BGS as well approximated by reductions in local genomic *N*_*e*_. Encouragingly, this is true even in spite of the imperfections of the B-map we used. PSMC, in contrast, produced biased estimates of effective population size (15% relative bias; Figure 1a), consistent with previous simulation studies [18, 19]. We next evaluated the bias of PSMC+ and PSMC under a simulation with BGS and changing effective population sizes. We simulated an *N*_*e*_(*t*) mirroring previous PSMC inferences on data from the YRI in 1KGP [25, 31]. PSMC+ was unbiased and PSMC underestimated the effective population size across the entire time range (Figure 1b).

**Figure 1.**
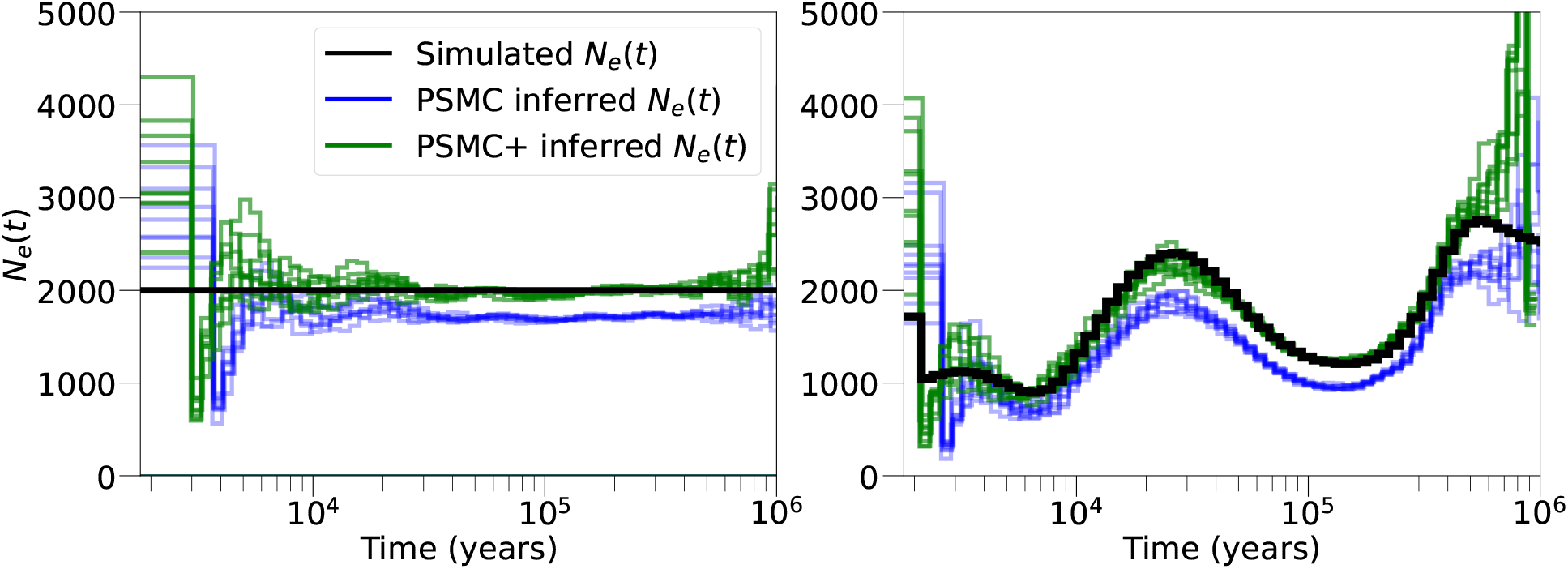
PSMC+ performance in forward-in-time simulations with widespread linked selection. **a)** Constant population size. **b)** Realistic demography, based on previous estimates of inferred *N*_*e*_(*t*) in West Africans [2, 30]. We simulated a demography (black line) and performed inference with regular PSMC (blue line), then PSMC+ (green line). The PSMC estimates of *N*_*e*_(*t*) are biased with a relative error of ∼ 15% throughout all time. Using a B-map allows PSMC+ to be approximately unbiased. We used the known map of simulated functional elements to create the B-map used given to PSMC+. 10 simulation replicates are shown for each evolutionary history.

PSMC+ leverages a model of local scaling of the effective population size to overcome the effect of linked selection on inferring *N*_*e*_(*t*). But even though the inferred *N*_*e*_(*t*) from PSMC is an underestimate, we find empirically that the general shape is accurate (Figure 1). This suggested to us the possibility that we can adjust for effects of BGS on PSMC’s inference of *N*_*e*_(*t*) by scaling the output according to the ratio of heterozygosity in exonic regions to the top 1% of regions most distant from an exon (Supplementary Figure 2). More concretely, PSMC works in units of *θ* and scales to absolute *N*_*e*_ and time in generations by using *µ*. If we scale the output with the ratio between *θ* calculated genome-wide and *θ* calculated in the top 1% of regions most distant from an exon, we can reduce the bias induced by BGS.

We evaluated the accuracy of PSMC’s inference of TMRCA in a simulation with and without BGS (Supplementary Figure 3). Both are biased, with loci that have true TMRCAs that are unusually ancient being consistently underestimated, and loci with true TRMCAs that are unusually recent being consistently overestimated – this is a simple regression-to-the-mean effect, which is expected when there is limited information at any locus [32, 33]. The level of bias in the simulation with BGS is slightly greater, though this effect is minimal. This indicates that even with pervasive BGS across the genome, the ability of PSMC to reconstruct the TMRCA is not strongly affected.

Finally, we evaluated the relative performances of PSMC+ and PSMC in the presence of mutation rate variation. If the mutation rate is not constant over the genome, this induces bias in PSMC inferred *N*_*e*_(*t*) which assumes rate homogeneity (Supplementary Figure 4a). If we feed PSMC+ the true mutation map, we are able to overcome these biases. While the true mutation map is unknown, it can be inferred from orthogonal measurements such as divergence per base pair between two distantly related species [34], the density of very rare mutations [35], or the nucleotide context [36, 37]. We note that the original PSMC paper suggested that the algorithm was robust to variation in mutation rate based on phylogenetic measures of mutation rate variation, but our results show that if the mutation rate variation is sufficiently large this is not the case. For example, drawing the mutation rate from a normal distribution with mean 1e-07 and variance 5e-08 produces significant bias in the inferred *N*_*e*_(*t*) (Supplementary Figure 4b). The advantage of feeding a mutation map into PSMC+ rather than simply relying on the PSMC algorithm is even greater for the ability to infer local TMRCAs. PSMC is not able to infer accurately the TMRCA across the genome in the presence of large variation in the mutation rate, but PSMC+ can recover this much more accurately given a mutation map (Supplementary Figure 4b).

### 4.2 Effect of background selection on autosome effective population size and coalescence time estimates

We studied the effect of background selection on the autosomes in humans. We binned the genome into 1MB segments and assigned those segments into five quantiles based on their average B-value in a high-resolution map of BGS in humans [26]. We then ran PSMC on each quantile separately (Figure 2; Methods). For the lowest B-quantile, we observe a reduction in effective population size of as much as 50% compared to the highest quantile. More generally, we confirm that the magnitude of effective population sizes scales with the strength of B-value, consistent with previously published findings. This is not an artifact of there being less heterozygosity in low B-value bins because the effect persists even when this parameter is fixed to the same value across bins (Supplementary Figure 5).

**Figure 2.**
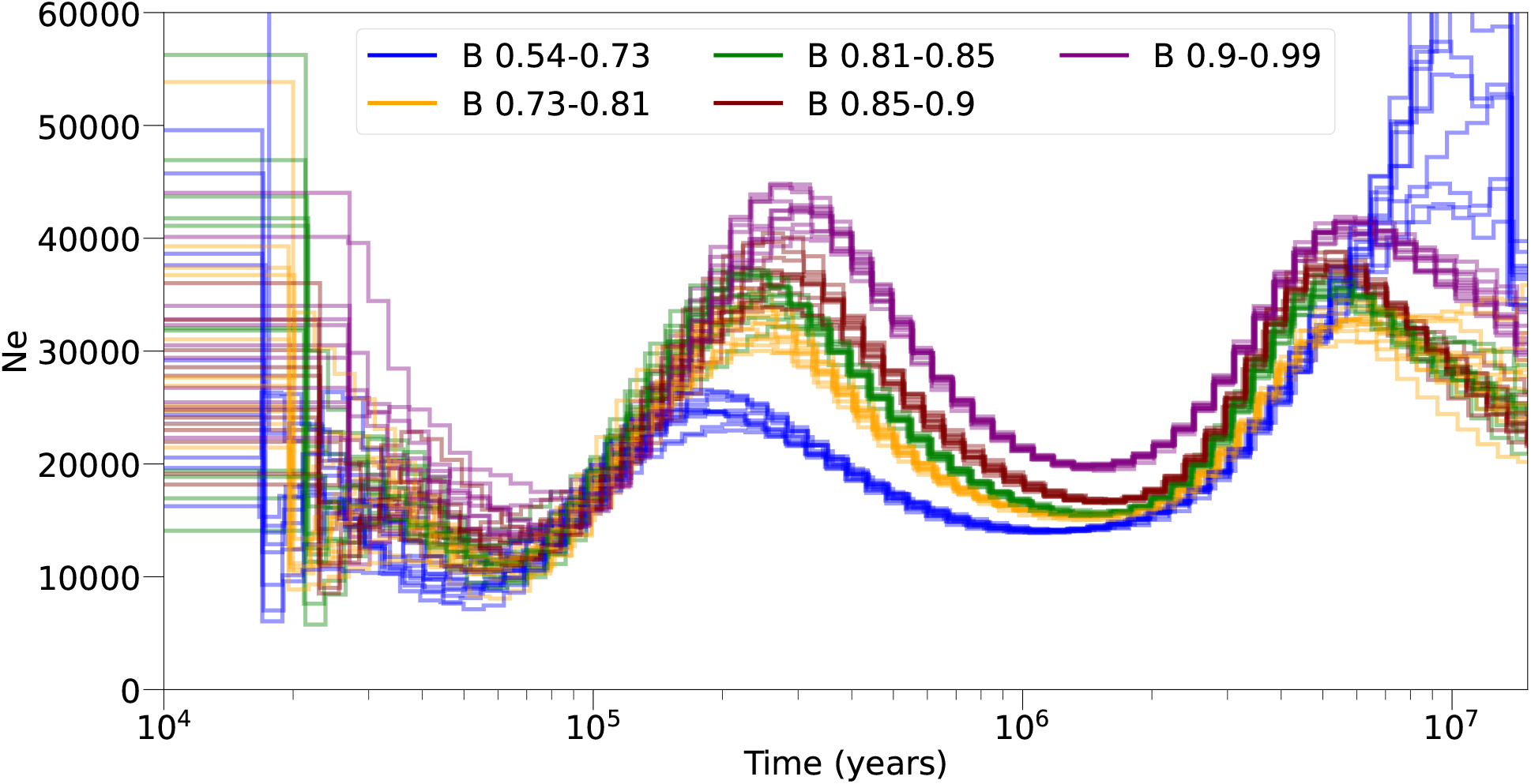
Inference of autosomal *N*_*e*_(*t*) in 80 YRI samples, in bins of b value. We binned the autosome into 5 equally sized bins based on the mean b value, and ran PSMC separately on each. We see substantial differences in the *N*_*e*_(*t*) curves across bins of B value, with the most neutral bin (B=0.90-0.99) showing the largest *N*_*e*_(*t*) across all time points.

We investigated the impact of BGS on inferred coalescent times. We examined the posterior decoding of the PSMC HMM, based on the inferred coalescent parameters for each YRI individual on the autosomes. At each position, we computed the posterior mean TMRCA for each YRI individual, and then took the maximum over all individuals. We computed the correlation between the maximum TMRCA and the B-map inferred by Murphy et al. and found a statistically significant Spearman correlation coefficient of 0.35 (p-value *<* 1*e* − 16). To check that this was not specific to method of PSMC, we used RELATE [38] and ARGweaver [39] to construct an ancestral recombination graph (see Methods), from which we extracted the maximum TMRCA across all YRI. The Spearman correlation between the maximum TMRCA and *b*-value was 0.34 (p-value *<* 1*e* − 16) and 0.19 (p-value *<* 1*e* − 16) in ARGweaver and Relate, respectively. We looked at PSMC’s inferred TMRCAs (for a diploid sample) and stratified these by quintiles of *b*-value. We observe that as the strength of BGS increases, the distribution of TMRCA gets younger (Supplementary Figure 6) (Kolmogorov-Smirnov test P-value *<* 1*e* − 16 for all pairwise comparisons). These observations are expected under the standard model that regions the genome that experience stronger BGS coalesce faster than neutral regions [22, 23].

We were curious whether we could leverage differences in how BGS would be expected to influence coalescence rates in a scenarios of panmictic size changes versus population structure, a well known identifiability problem in PSMC [40, 41]. We performed two sets of simulations with background selection: one with ancestral population structure and one with changes in the *N*_*e*_(*t*), where the size changes are set such that each simulation has the same coalescence rate (Supplementary Figure 7). We stratified the simulated genomes according to the amount of background selection they experienced, ran PSMC, and found that the profiles were very similar (Supplementary Figure 7). This suggests that stratifying by intensity of expected background selection may not be useful for distinguishing ancestral population structure from changes in effective population size.

### 4.3 Application of PSMC+ to human demographic history

We next applied PSMC+ to high-coverage whole genome sequencing data from YRI sequenced by the 1000 Genomes Consortium (total n = 80 diploid individuals; Methods). We used the B-map inferred by Murphy et al. and observe that the inferred *N*_*e*_(*t*) by PSMC+ is elevated with respect to the PSMC inference (Figure 3, green and blue lines respectively), consistent with the results on simulations (Figure 1). Using a simple B-map calculated based on distance to exon or simply scaling the PSMC inference achieves a similar result to the PSMC inference (Figure 3, pink and yellow lines respectively), verifying the utility of these approaches.

**Figure 3.**
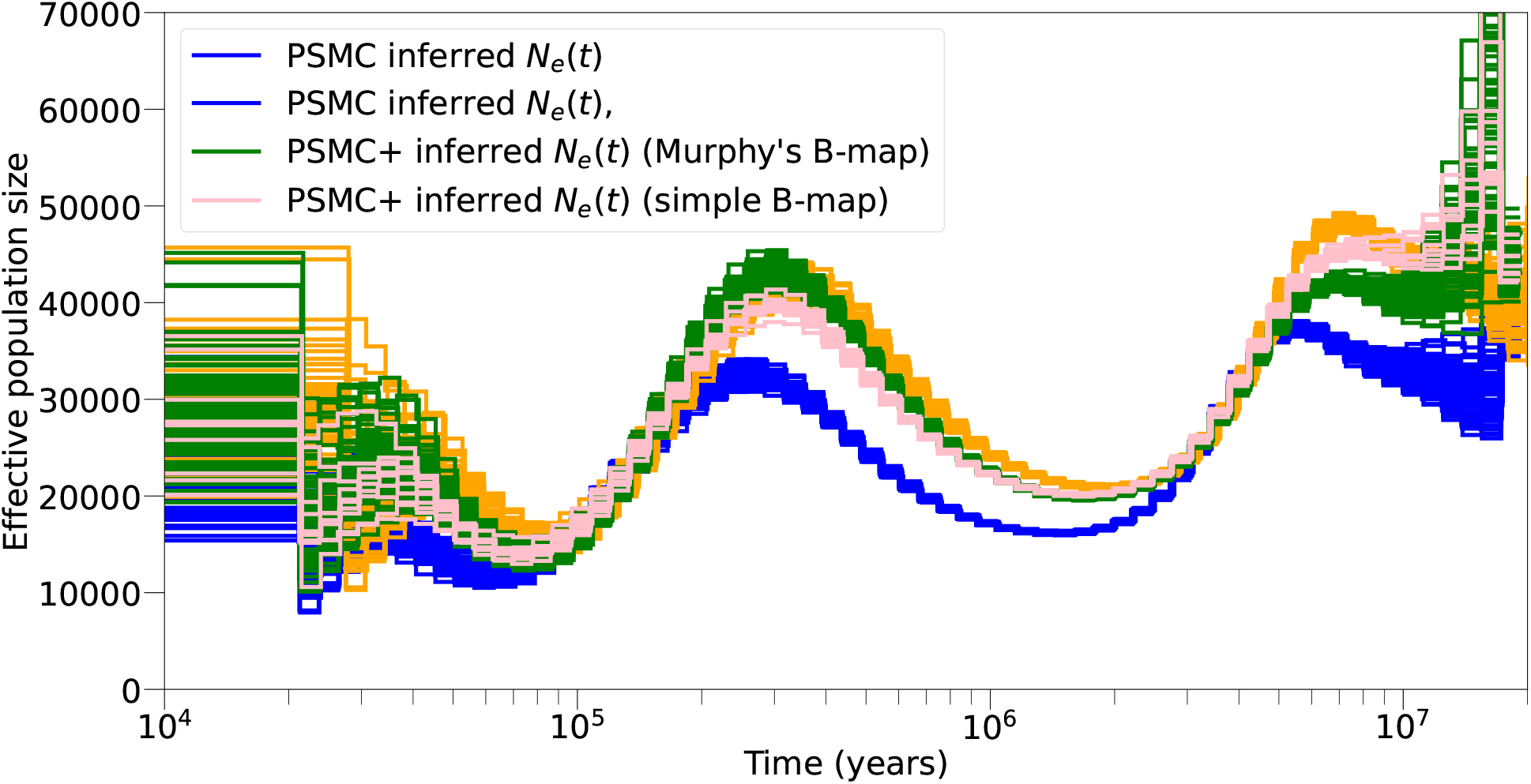
Inference of *N*_*e*_(*t*) on 80 YRI samples. Inference from natural PSMC is shown in blue, PSMC+ with Murphy’s B-map in green, PSMC+ with a simple B-map constructed based on distance to exon in pink, and PSMC inference scaled by the ratio of exonic to non-exonic heterozygosity in gold.

### 4.4 Effect of background selection on X to autosome effective population size ratio estimation across primate species

We studied the ratio of X chromosome to autosome *N*_*e*_(*t*) through time (*R*(*t*)) on numerous great ape species, without applying any correction for background selection since we did not have B-maps constructed in the same way on both the autosomes and the X chromosome, and since we did not have B-maps in non-humans. We hypothesized *R*(*t*) would deviate from 3/4 due to the more extreme effects of linked selection on chromosome X compared to the autosomes [6]. We ran PSMC on the X chromosome and autosomes separately for humans, chimpanzees, gorillas, and orangutans (Supplementary Figure 8).

We observe strong deviations from the expected 3/4 ratio for all species (Figure 4). A basic coalescent model with no selection and constant size suggests an upper and lower bound for *R*(*t*) of 9/8 and 9/16, respectively (see Methods), although it has been demonstrated that extremely strong founder events can generate *R*(*t*) *<* 0.3 [42]. In chimpanzees, gorillas, and orangutans, more recently than 100ky we observe that *R*(*t*) is below 9/16. In humans we observe a similar effect, though between 100ky to 300ky. More recently than ∼100ky we observe *R*(*t*) is elevated above 9/8. Interestingly, *R*(*t*) is around 3/4 in all species at 1My, which is the expectation under neutrality.

**Figure 4.**
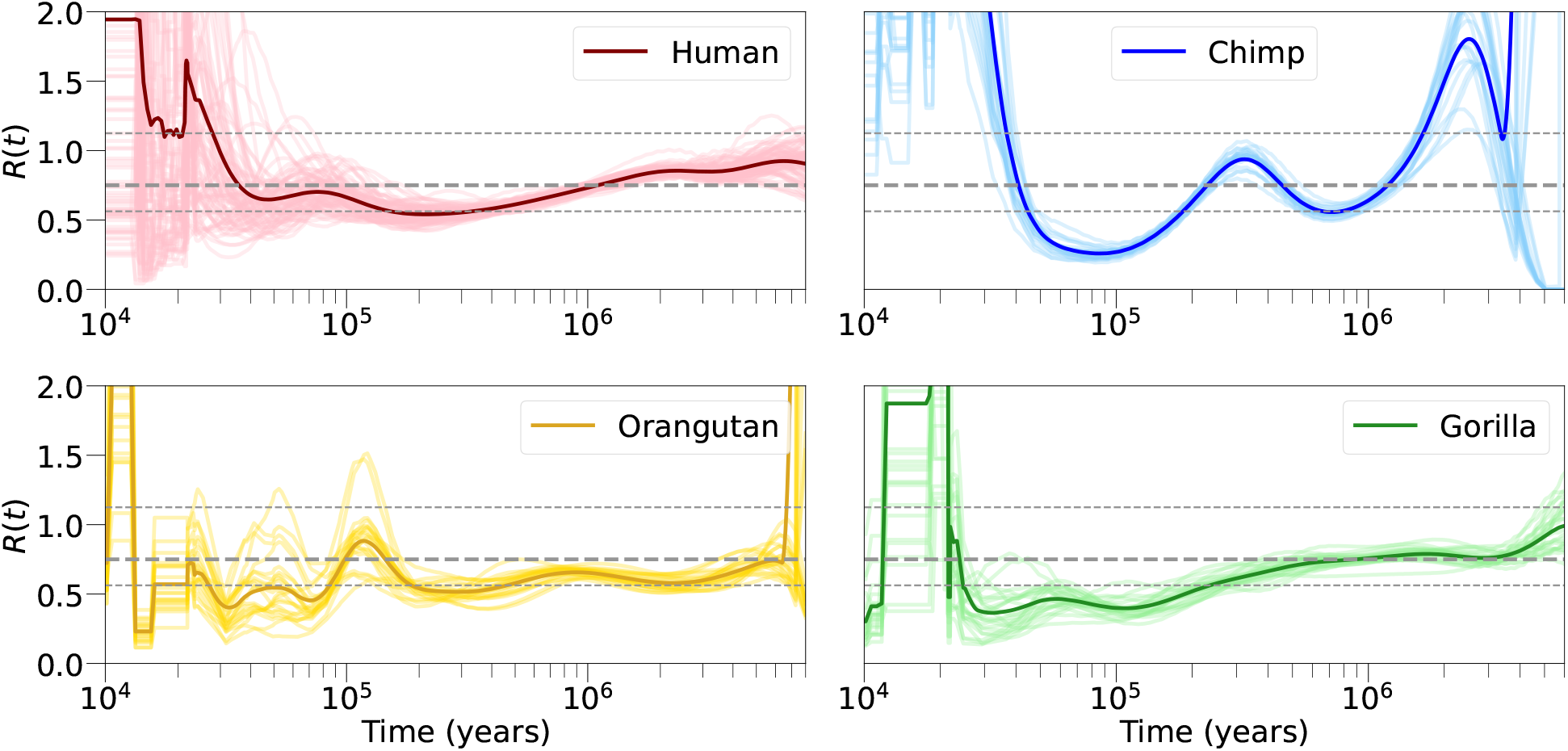
The ratio of *N*_*e*_(*t*) inferred on the X chromosome to the autosomes, for humans, chimpanzees, gorillas, and orangutans. The mean is shown in the darker line, which is calculated across all individuals and bootstraps as shown in the lighter line. Under neutrality we would expect this value to be 0.75 (thick, gray, dashed line), but we observe strong deviations from this in each species, often exceeding the theoretical bounds (thin, gray, dashed lines).

In each species, PSMC does not detect a founder event as severe as required by Pool and Nielsen [42] to give *R*(*t*) *<* 0.3, so variations in *R*(*t*) are likely not attributable to changes in *N*_*e*_(*t*) alone. A possible explanation for the extreme values of *R*(*t*) is uncorrected background selection, although we cannot rule out additional contributions to the signals such as changes in life history traits or mutation rate variation [43, 44]. To test the hypothesis that background selection drives changes in the ratio over time, we computed the ratio of effective population sizes of the lowest B-value bin to the highest B-value bin as computed on the human autosomes (Supplementary Figure 9). Similar to the X:A ratio, we observe changes in the ratio over time, suggesting that uncorrected effects even here may be driving the signal. As discussed above, it is not clear how to run PSMC+ on chromosome X because Murphy et al. did not release a map of background selection on chromosome X, and even if it was constructed, it would be impossible to be confident it had the same resolution as the autosomal one. We also did not run PSMC+ on the other great ape species because high-resolution maps of background selection do not yet exist for those species.

## 5 Discussion

We developed a new method, PSMC+, for estimating effective population size trajectories (*N*_*e*_(*t*)), that can incorporate local variation in coalescent or mutation rates along the genome. Simulations indicate our method is unbiased in the presence of background selection (BGS). By applying PSMC+ to data from the 1000 Genomes project, we identify as much as a ∼30% increase in the effective population size around 300kya relative to previous estimates. We also study how autosomal *N*_*e*_(*t*) changes on the autosomes as a function of BGS, and find that at ∼400ky in the strongest *b*-value bin *N*_*e*_(*t*) is 50% the size of *N*_*e*_(*t*) in the weakest *b*-value bin. This qualitatively matches comparisons of the X chromosome to the autosomes where a similar maximum is achieved at this time point, suggesting background selection may be an important factor in shaping the coalescent trajectory of the X chromosome. We also test our method on a low-resolution map of BGS, the construction of which does not require high quality annotations of fitness effects. This approach seems to work well and we anticipate it will be useful in a wide variety of non-model organisms. Finally, we test the accuracy of coalescence time inference in the presence of BGS and find that the PSMC posterior decoding is robust.

Our study has several implications for future analyses. First, while comparisons of X chromosome to autosome effective population sizes can be informative of life history traits [45], our results indicate that differences in the strength of background selection is an important factor in shaping effective population size, which complicates interpretations of X to autosome comparisons. Second, our results indicate that background selection can have a substantial impact on demographic inference. Most current methods do not explicitly handle background selection, which leads to bias in the parameter estimates. Because of differences in background selection across species, this is expected to affect some species more than others [8]. Third, simulations suggest that background selection does not reduce the accuracy in PSMC’s inferred coalescence times, even though this is ignored in the underlying model. This is important for other ARG inference methods [39, 38, 46, 47, 48, 49] which also ignore the effects of selection on genealogies. Our claims that background selection affects inference of *N*_*e*_(*t*) but does not affect inference of coalescence times may seem contradictory. We argue that, from a simulation-based perspective, the true effective population size (defined as the inverse of the coalescence rate) really is smaller than the effective population size as specified in the simulation, because deleterious mutations have purged haplotypes from the population. Thus the quantity we are reconstructing with PSMC+ can be thought of as the “neutral effective population size”, which we define as the what the effective population size would be in the absence of deleterious mutations.

We highlight several areas of future work. First, it may be possible to adjust the transition probabilities of PSMC’s HMM rather than the emission model as we do here, which would more faithfully model the perturbations in genealogy due to purifying selection [50]. Second, future work will illuminate the effects of BGS on the human X chromosome, as well as the autosomes for other species. This will allow greater resolution in how BGS affects *R*(*t*). Third, numerous maps of the *de-novo* mutation rate across the genome exist for humans [35, 36, 37, 51, 52], and studying how these affect inference of *N*_*e*_(*t*) in PSMC+ is a possible further direction.

Our study has several limitations. First, we follow Nordborg et al. [23] and Hudson and Kaplan [22] in modeling background selection as a reduction in the local effective population size. While this has been shown to capture broad-scale effects of background selection, it assumes new mutations are so deleterious that they cannot fix in a population. Our simulations indicate that utilising this model reduces bias in the inferred *N*_*e*_(*t*), but it will not capture all the effects of BGS [53] as the model does not capture weak selection or the dynamics of rare variants. However, as we focus on *N*_*e*_(*t*) more anciently than ∼10kya, we do not expect weak selection or rare variants to be important for the inference we perform here. A possibility for future work would be to study how newer models of BGS that explicitly capture weak selection [54] affect more recent estimates of *N*_*e*_(*t*), which in principle could be studied by feeding PSMC+ more chromosomes and using a composite likelihood approach as in MSMC. Despite these limitations, our study provides an improved understanding of the effects of background selection on demographic history and highlights the need to consider the impact of non-neutral forces in demographic inference.

## 6 Methods

### 6.1 Data collection, processing, and annotations

We downloaded aligned reads (BAM files) from 81 female individuals from the 1000 Genomes Phase 3 high coverage sequencing release from the New York Genome Center [31]. These files are aligned to the version hg38 of the human reference genome. We called SNPs with bcftools mpileup, and set minimum mapping quality 20, minimum base quality 20, and adjusted mapping quality 50. We called SNPs with bcftools call and then masked rejoins of the genome where the coverage was less than half or more than double the mean coverage from chromosome 20. We also masked regions of the genome according to a strict mappability mask for hg20. One individual had excessively low heterozygosity, likely reflecting sequencing or bioinformatic errors, so was not used in subsequent analysis.

The procedure for processing the other great apes was very similar. We downloaded aligned reads (BAM files) from EBI [55], including 4 chimpanzees (2 female), 6 gorillas (4 female), and 3 orangutans (1 female). These were aligned to their own reference genomeS: Pan tro 3.0 (UCSC: panTro5); gorilla, Gor-Gor4.1 (UCSC: gorGor4); orangutan WUGSC2.0.2 (UCSC: ponAbe2). We built a mappability mask for each reference genome using Heng Li’s SNPable.

We downloaded a map of background selection released by Murphy et al. We converted their files to bed format and used LiftOver [56] to convert the coordinates from hg19 to hg20 (GRCh37 to GRCh38). We filled in missing values b values with 1, which largely overlapped with the uncallable regions from the hg20 mappability mask.

### 6.2 Scaling coalescent time

In humans, we use a mutation rate per generation per base pair of 1.25e-08 [57, 58, 59, 60] and generation time of 29 years [61]. As suggested in [62], we use the following mutation rate, generation time parameters in chimpanzees 1.78e-08, 24, gorillas 1.42e-08, 19, orangutans 2.03e-08, 27.

### 6.3 Stratifying the autosome by B value and running PSMC

We stratified the autosome into 5 equally sized bins of b value (measured by the amount of autosome in each bin). The cutoffs b value cutoffs we used were: [0.53-0.72), [0.72-0.8), [0.8-0.85), [0.85-0.89), [0.89-0.98]. The B value varies continuously along the genome. This is a problem for our analysis, because we would like to model local genealogies, which extend over a stretch of the genome. To solve this, we averaged the B value over 1Mb. Then, we treated each 1Mb segment as a separate chromosome, and ran PSMC separately for each B value bin. Our choice of 1Mb is motivated by the fact that the B value does not vary much across segments of 1Mb (average standard deviation of B value across 1Mb segment, dragged in windows of 100kb = 0.065). We did not analyze the X chromosome because Murphy et al., 2023 did not release a B value map for the X chromosome.

### 6.4 Adjusting the PSMC model to account for heterogeneous rates

PSMC [2] is a HMM where the hidden states are the discretised coalescence times *Z* = (*z*_1_, …, *Z*_*L*_) and the observations *X* = (*x*_1_, …, *x*_*L*_) are the series of homozygotes or heterozygotes along a diploid chromosome. The transitions between the hidden states are governed by the SMC or SMC’ framework, and are a function of the population size changes. The emission model describes the probability of a mutation arising given a coalescence time. PSMC works in units of the population mutation rate, *θ* = 4*Neµ*, where *N*_*e*_ is the long term effective population size and *µ* is the de-novo mutation rate per generation per base pair. In our implementation, the genome is binned into *k* base pairs (typically *k* = 100) and thus *x*_*i*_ takes values in (1, …, *k*). The number of mutations in a bin is then modeled as a Poisson and we write:

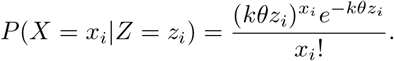

Given a map of variations in *θ, F* = (*f*_1_, …, *f*_*L*_), the emission probabilities can be simply adjusted with:

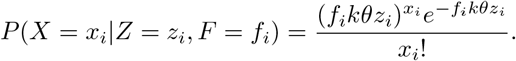

These can then easily be built into the PSMC model with standard HMM machinery [63] [64] [65].

### 6.5 Simulations of background selection

We performed forward in time simulations using SLiM v3.7.1 [29]. We simulated 150 megabase chromosomes, which are comparable in size to human chromosome 8 and X. We rescaled the population sizes and mutation rates by a factor of 10 to ensure our simulations did not consume an impractical amount of memory. We simulated exons using a realistic map. We simulated non-synonymous mutations with selection coefficients from from a gamma-distributed DFE with shape=0.513 and scale=5.38, based off of Kim et al. [66]. We simulated a burn in of 20,000-40,000 generations with a population size of 2,000 diploid individuals.

We simulated 20 individuals with 10 replicates. We used a mutation rate of 1.25e-7 per base pair per generation, which is ∼10x higher than the human mutation rate [67], because we wanted the genome-wide heterozygosity to be of similar magnitude to that in humans. We used a constant recombination rate of 1e-8 per basepair per generation. We did not scale the recombination rate as we found doing so reduced the strength of BGS due to linkage being too weak. Our simulations recapitulate to a large extent the effect of BGS as seen in humans, when measuring diversity as a function of distance from non-synonymous mutations (Supplementary Figure 5).

### 6.6 Theoretical bounds of *R*(*t*) under neutrality

Here we derive the upper and lower bounds of *R*(*t*) - defined as *N*_*e*_(*t*) inferred on the X chromosome divided by *N*_*e*_(*t*) inferred on the autosomes - for a panmictic population of constant size. Suppose we have *m* reproducing males and *f* females. Then the probability that two random contemporaneous children share a father is 1*/m* and 1*/f* for the mother. Consider two uncoalesced autosomal lineages; the probability that in a given generation they both go through a female is 1/4 and similarly for a male. Then, the coalescent rate on the autosomes is

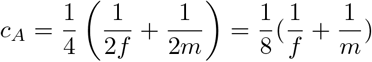

Similarly, for two uncoalesced X chromosomes lineages, the probability they both go through a female is 4*/*9, through a male is 1*/*9, so the coalescent rate is

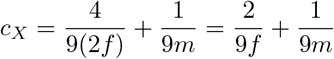

So

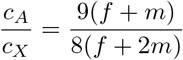

For *f >> m* we get 9/16, and for *m >> f* we get 9/8, so we obtain bounds:

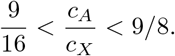

We note that dramatic changes in population size or population structure can create more extreme ratios of *R*(*t*) [68] [42].

### 6.7 Constructing a simple Bmap

Under the expectation that parts of the genome most distant from coding regions are least likely to be affected by linked selection, we calculated a simple B-map, by first computing the normalized distance of each base pair to its closest exon (measured in genetic distance). As the effects of BGS are better modeled at length scales larger than one base pair, we take the mean exon distance in a window of some size w. Moreover, because the likelihood of a recombination event between two loci decreases exponentially as the physical or genetic distance between them increases, we can create a new B-map by simply transforming the normalized distance *x* with

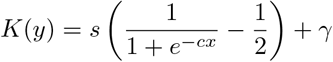

where *γ* controls the minimum *b* value, *c* controls the rate of decay, and *s* is defined such that *K*(*y*) is always between *b* and 1:

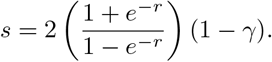

In both our simulations (Figure 1) and the YRI (Figure 3), we set *w*=1e+06,*γ* = 0.6 and *c* = 100.

### 6.8 Software and data availability

PSMC+ is freely available to download and use github.com/trevorcousins/PSMCplus.

## 7 Acknowledgements

We gratefully acknowledge the fruitful discussions with members of the Reich laboratory and the Durbin group. This work was supported by the National Institutes of Health grant HG012287 (D.R.), by the John Templeton Foundation grant 61220 (D.R.), by the Howard Hughes Medical Institute (D.R.).

## 8 Supplementary Figures

**Supplementary Figure 1:**
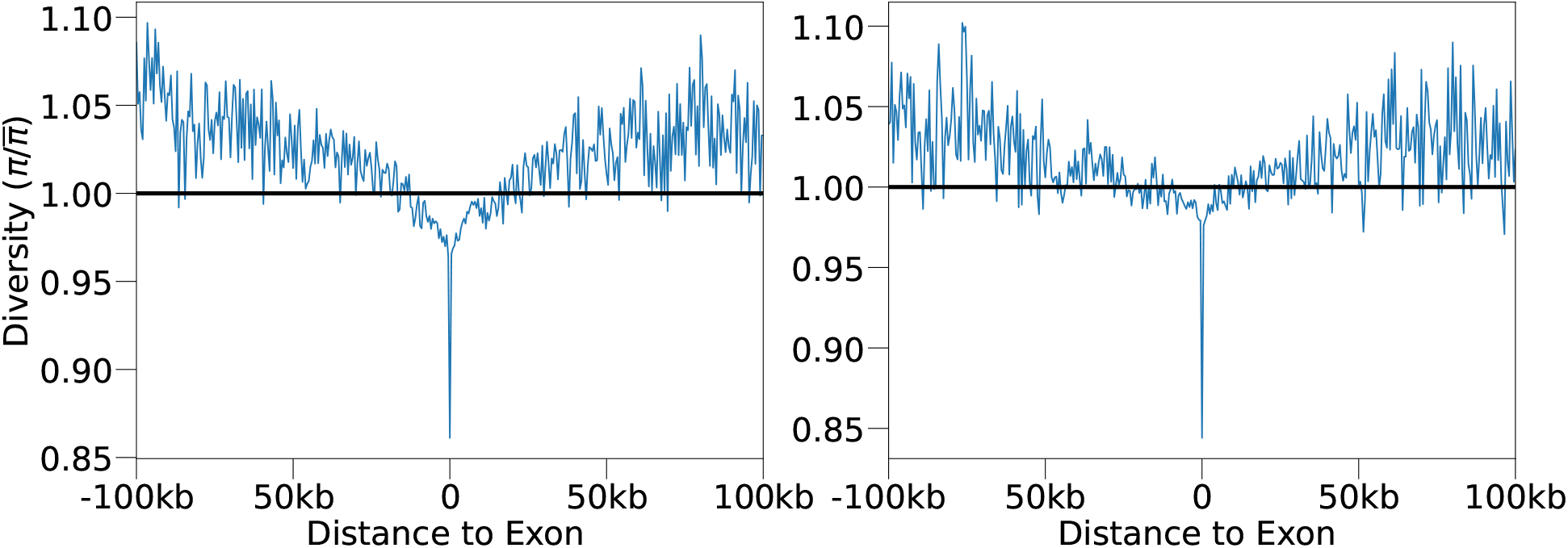
Our forward-in-time simulations of background selection show a similar effect in shaping genome wide diversity as seen in humans [26] We calculate the observed diversity as a function of distance to exon, and see a strong positive association for a) the constant-sized population and b) the changing-size population. We calculate diversity relative to the genome wide average, as indicated by the black line.

**Supplementary Figure 2:**
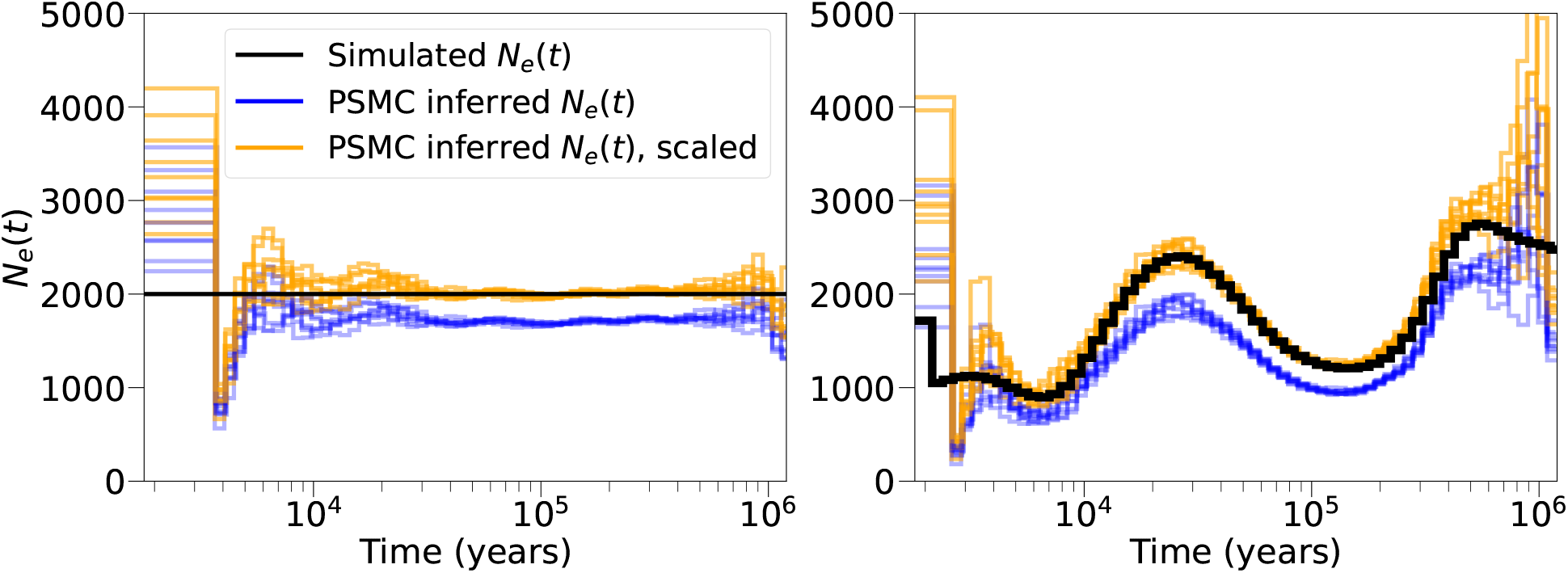
PSMC’s inference of *N*_*e*_(*t*) is biased in simulations with widespread linked selection (blue lines). Scaling PSMC’s inference of *N*_*e*_(*t*) (gold lines) is able to accurately overcome these effects.Same simulations as in Figure 1: **a)** Constant population size. **b)** Realistic demography, based on previous estimates of inferred *N*_*e*_(*t*) in West Africans [2, 30].

**Supplementary Figure 3:**
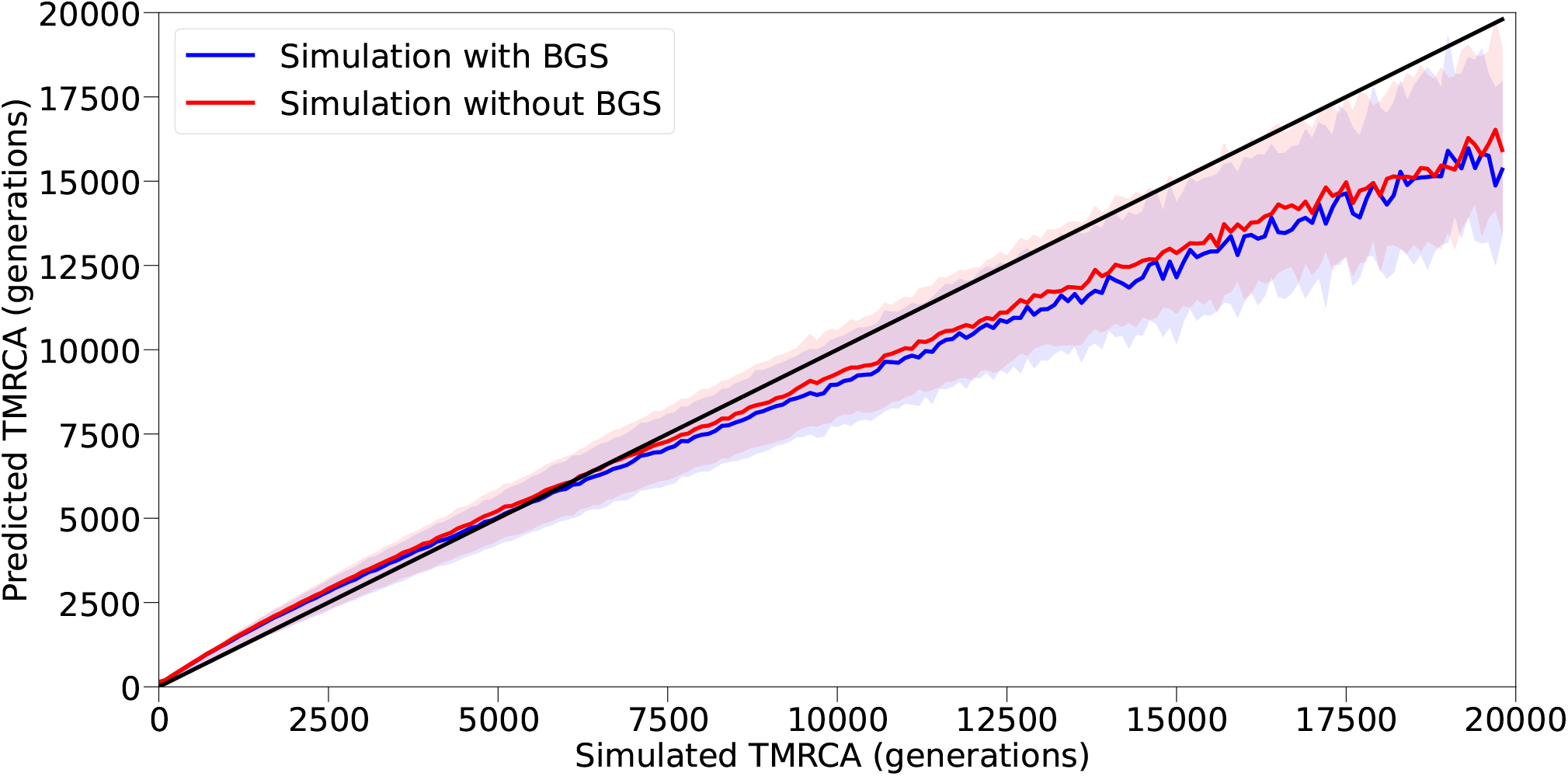
Accuracy in the inferred coalescence times from PSMC, in a simulation with and without BGS. The blue line shows inference in a model with BGS, and the red line without. We calculated accuracy by taking the posterior mean at each position and comparing this to the simulated value.

**Supplementary Figure 4:**
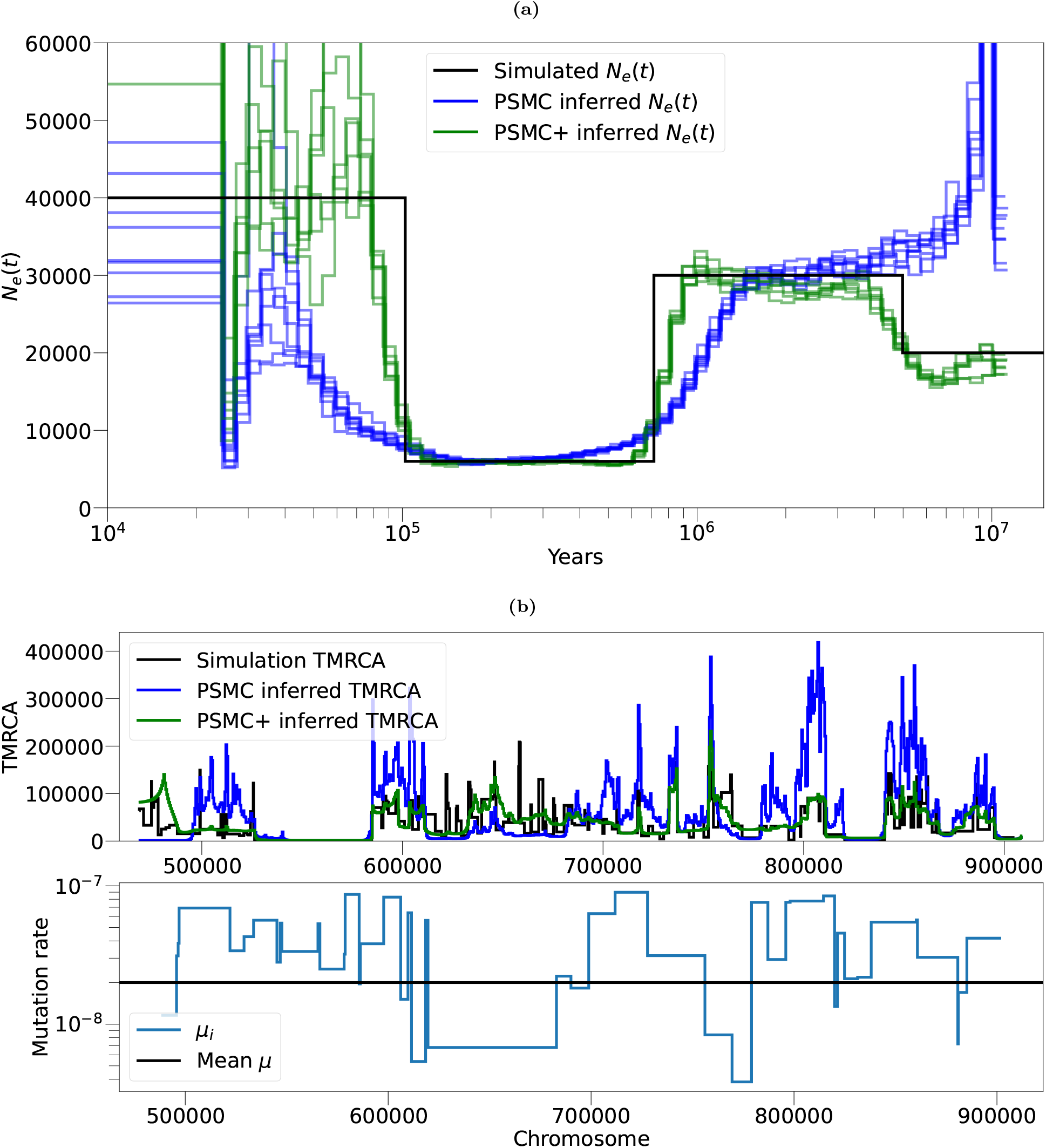
The effect of mutation rate variation in inferring *N*_*e*_(*t*) or the TMRCAs across the genome. An arbitrary mutation rate map M that changes every d base pairs was generated, where d is exponentially distributed with rate 100kb. The mutation rate in each interval was drawn by taking the absolute value from a normal distribution with mean 1e-07 and standard deviation 5e-08. The recombination rate was set as 1e-08. a) Inferring *N*_*e*_(*t*) with PSMC or PSMC+. The simulated *N*_*e*_(*t*) is shown in black. PSMC inference on a simulation with mutations generated by M shown in blue, which is not able to accurately recover *N*_*e*_(*t*). PSMC+ inference on the same simulation, is able to overcome the effect and infer *N*_*e*_(*t*) more accurately. b) Inference of TMRCA across the genome (top panel). The black line represents the simulated TMRCA. Posterior mean from the PSMC decoding is shown in blue, which is not able to capture the TMRCA distribution. Posterior mean from the PSMC+ decoding is shown in green, which is able to overcome the effect and infer the TMRCA more accurately. Bottom panel shows the local variations in M.

**Supplementary Figure 5:**
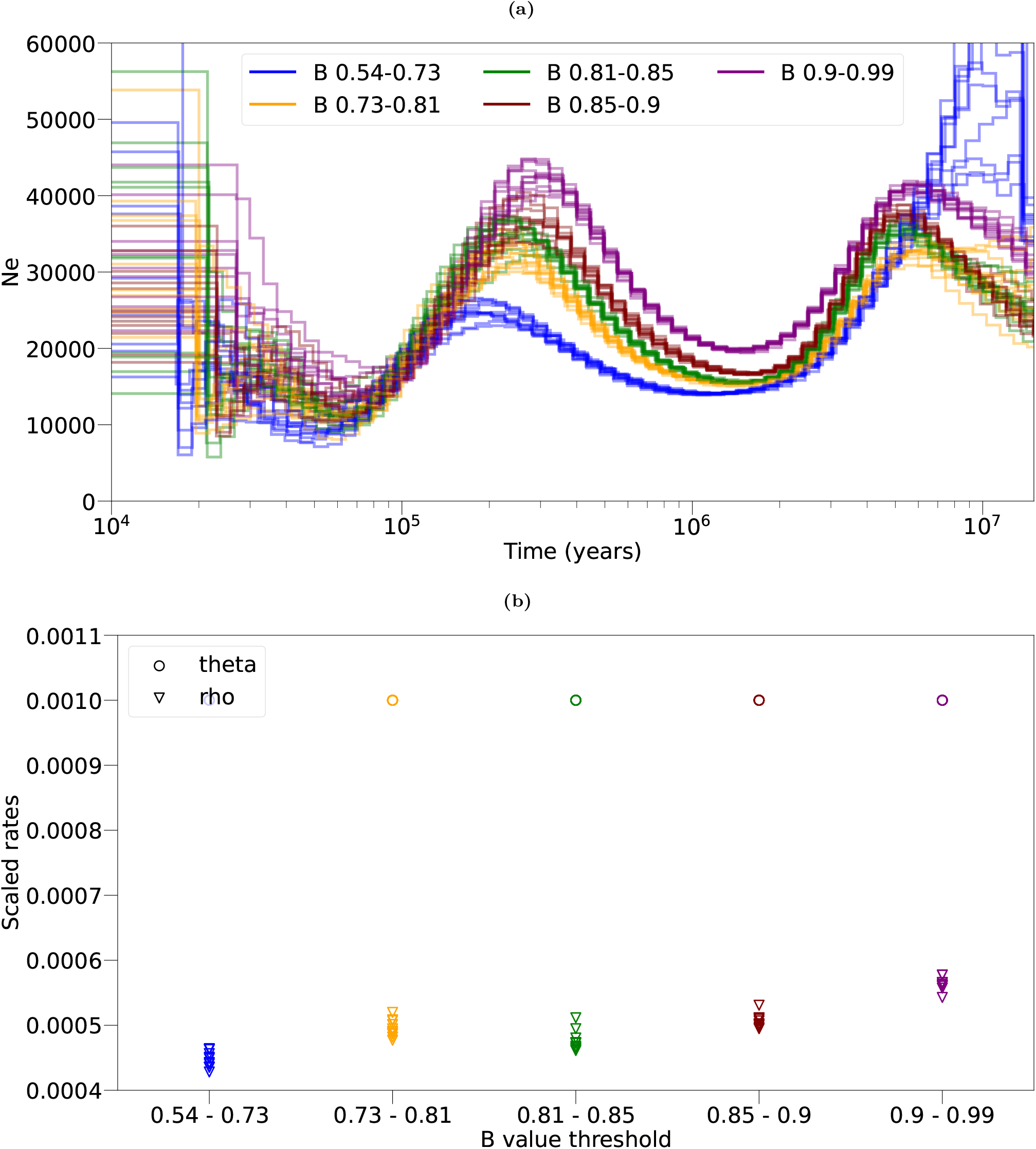
a) Inference of autosomal *N*_*e*_(*t*) in 80 YRI samples in bins of b value, though with fixed to the same value across each bin (similar to Figure 2 which used a calculated from the data). b) Each b quintile uses =0.001 (circles), but rho (triangles) is inferred as part of the EM algorithm and varies per quintile. This demonstrates that the differences in inferred *N*_*e*_(*t*) per b are not attributable to difference in heterozygosity per quintile.

**Supplementary Figure 6:**
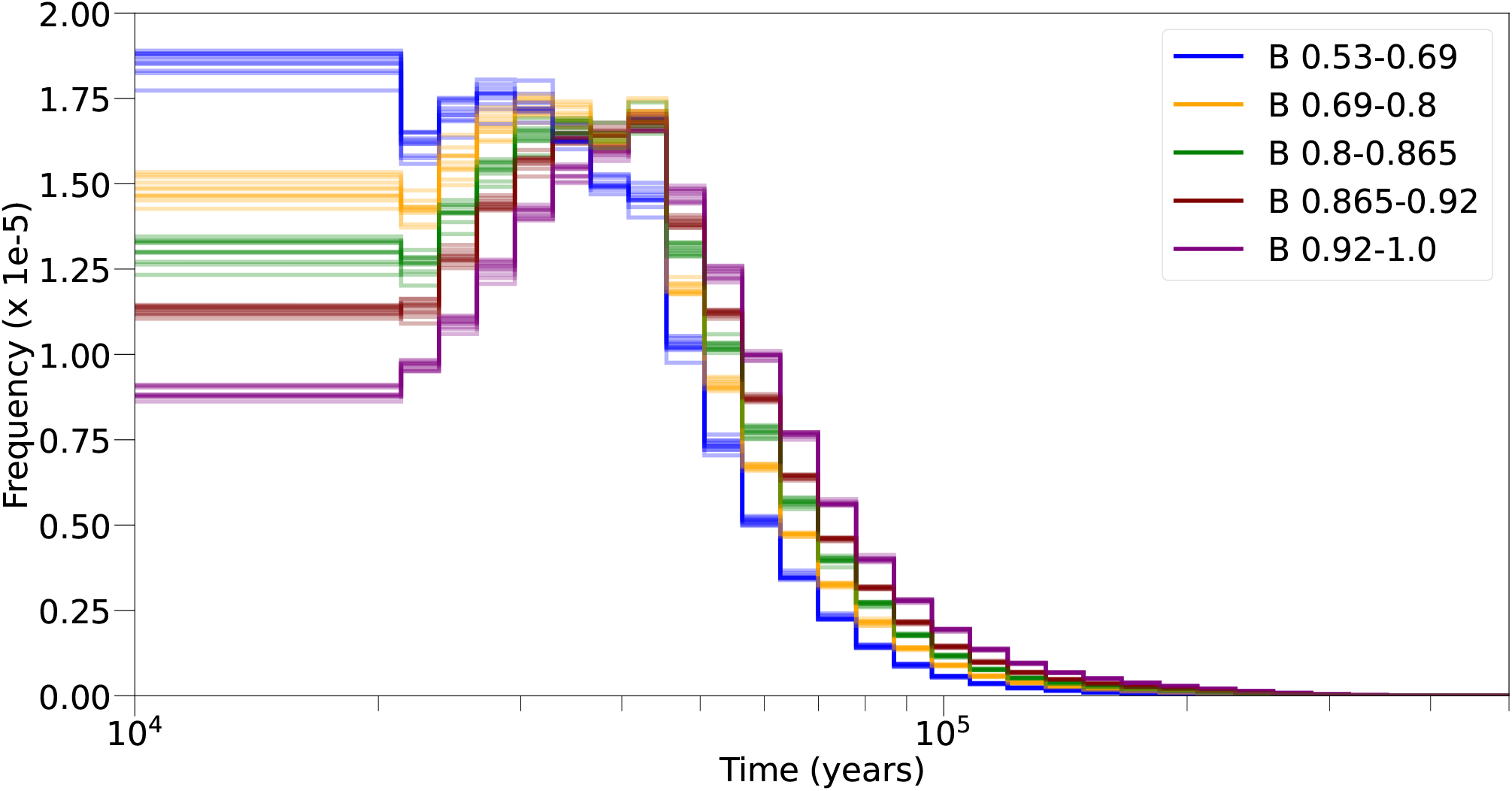
Effect of background selection on inferred TMRCA. We plot the distribution of TMRCA in coalescent units in quintiles of *b* value. In the lowest bin of *b* value, we see an excess of recent pairwise TMRCA, consistent with the action of linked negative selection lowering the effective population size and reducing the TMRCA.

**Supplementary Figure 7:**
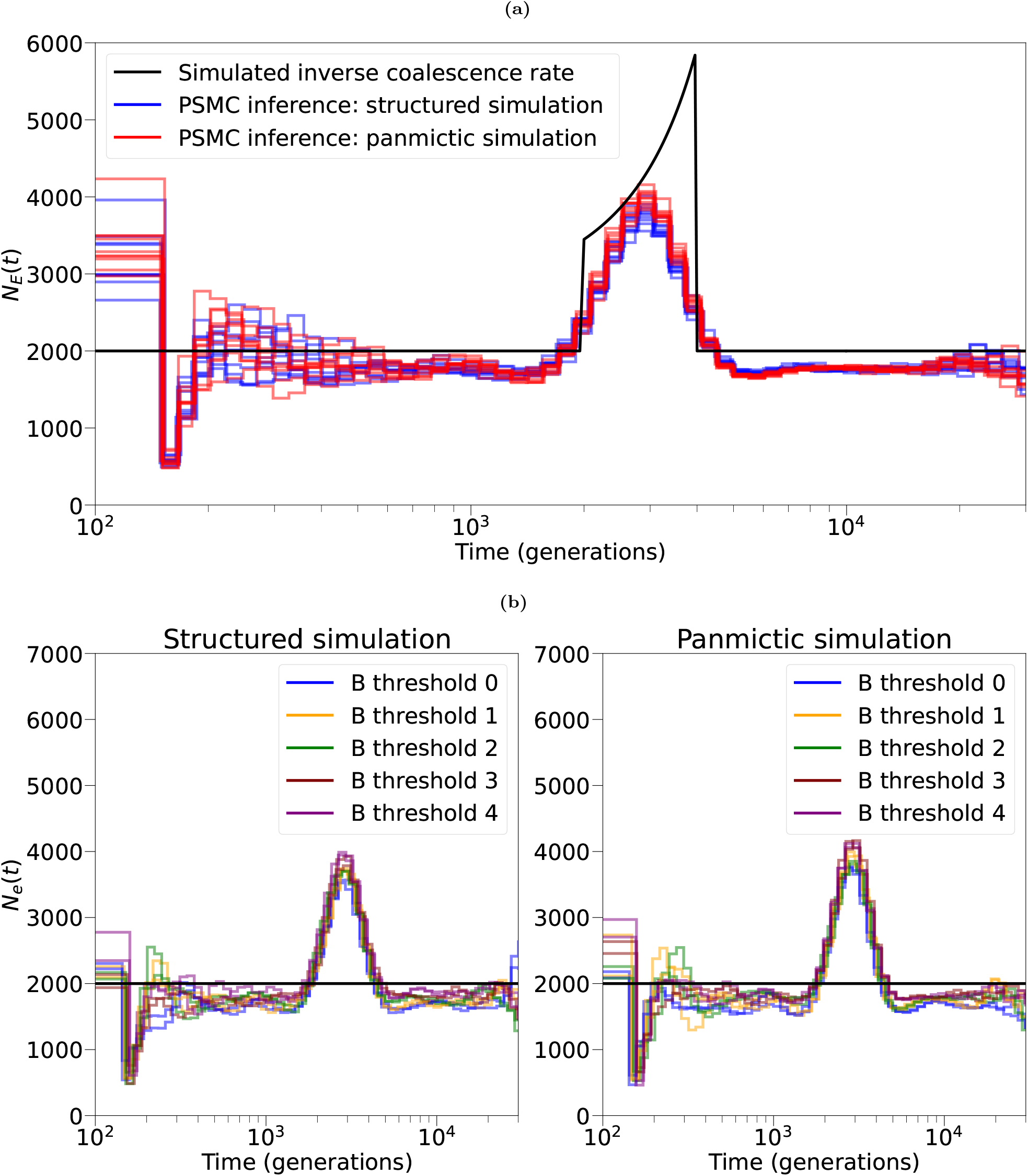
Inference of *N*_*e*_(*t*) on BGS simulations, where the evolutionary history is either panmictic or structured. The structured simulation has constant population size and a 30% admixture fraction; the panmictic simulation has changes in the effective size such that the coalescence rate matches the structured simulations. a) The simulated inverse coalescence rate is shown in the black line; PSMC’s inferred *N*_*e*_(*t*) on the structured simulation is shown in blue, and the panmictic in red. b) We stratified the genome into 5 quintiles based on the strength of BGS, then ran PSMC for both the structured simulation (left) and panmictic simulation (right).

**Supplementary Figure 8:**
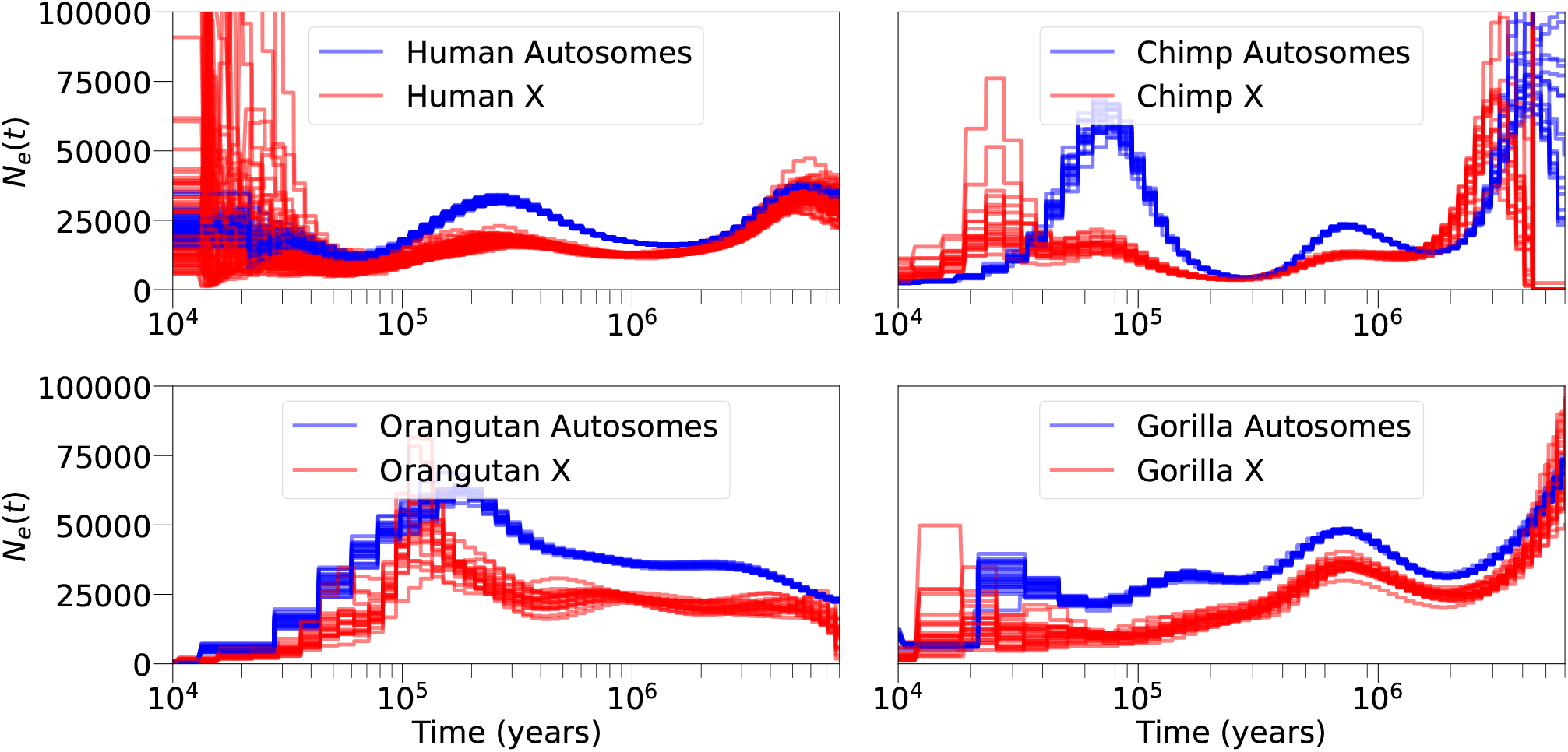
PSMC’s inferred *N*_*e*_(*t*) on the autosomes and X chromosome humans, chimps, gorillas, and orangutans.

**Supplementary Figure 9:**
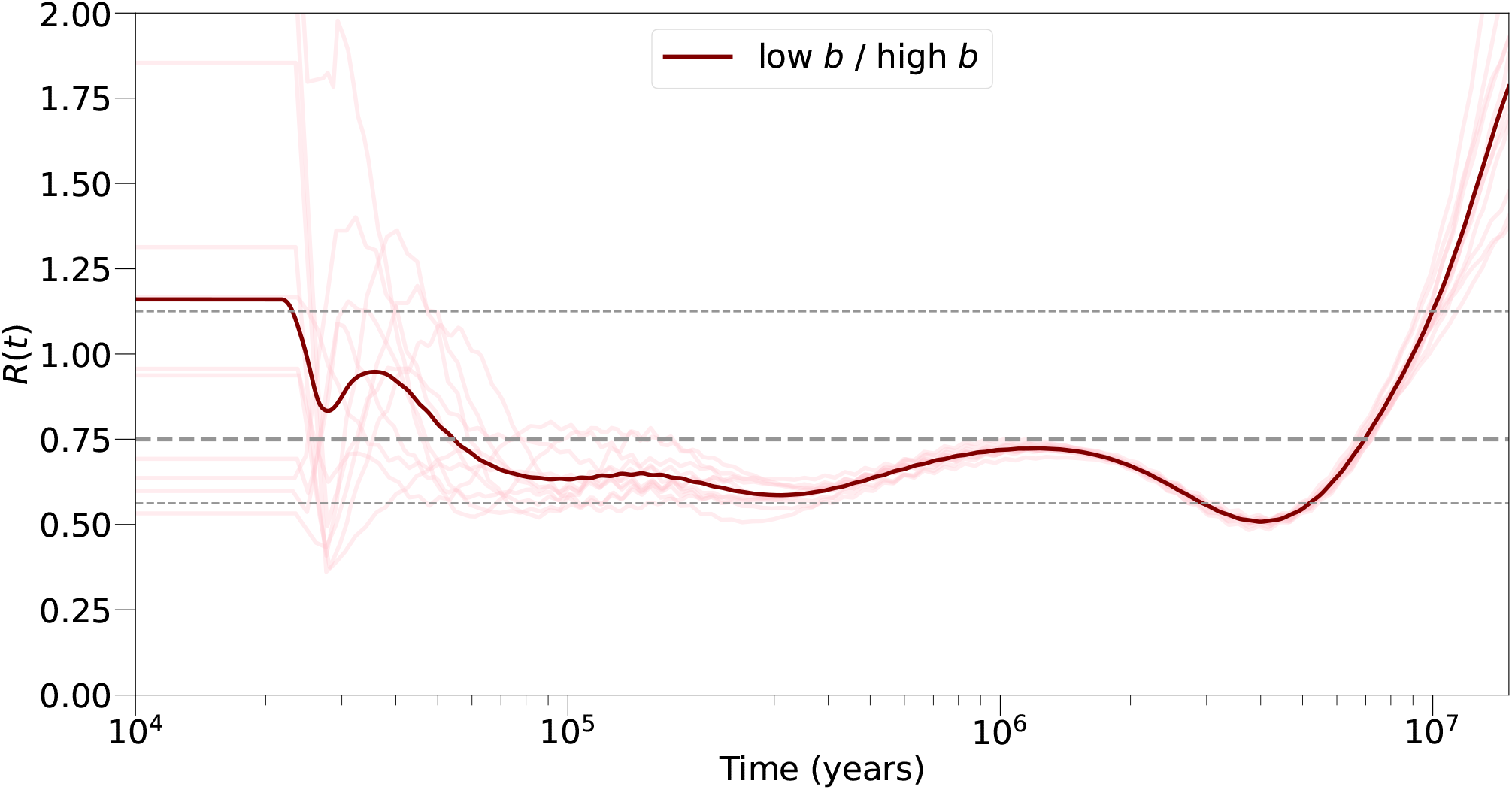
The ratio of *N*_*e*_(*t*) inferred on the strongest b value bin to the weakest b value bin, in Figure 2 (blue and purple lines, respectively).

## References

[1] Brian Charlesworth. Effective population size and patterns of molecular evolution and variation. Nature Reviews Genetics, 10(3):195–205, 2009.

[2] Heng Li and Richard Durbin. Inference of human population history from individual whole-genome sequences. Nature, 475(7357):493–496, 2011.

[3] Stephan Schiffels and Richard Durbin. Inferring human population size and separation history from multiple genome sequences. Nature genetics, 46(8):919–925, 2014.

[4] Sara Sheehan, Kelley Harris, and Yun S Song. Estimating variable effective population sizes from multiple genomes: a sequentially markov conditional sampling distribution approach. Genetics, 194(3):647–662, 2013.

[5] Jeffrey P Spence, Matthias Steinrücken, Jonathan Terhorst, and Yun S Song. Inference of population history using coalescent hmms: review and outlook. Current opinion in genetics & development, 53:70–76, 2018.

[6] Graham McVicker, David Gordon, Colleen Davis, and Phil Green. Widespread genomic signatures of natural selection in hominid evolution. PLoS genetics, 5(5):e1000471, 2009.

[7] Ryan D Hernandez, Joanna L Kelley, Eyal Elyashiv, S Cord Melton, Adam Auton, Gilean McVean, 1000 Genomes Project, Guy Sella, and Molly Przeworski. Classic selective sweeps were rare in recent human evolution. science, 331(6019):920–924, 2011.

[8] Russell B Corbett-Detig, Daniel L Hartl, and Timothy B Sackton. Natural selection constrains neutral diversity across a wide range of species. PLoS biology, 13(4):e1002112, 2015.

[9] Eyal Elyashiv, Shmuel Sattath, Tina T Hu, Alon Strutsovsky, Graham McVicker, Peter Andolfatto, Graham Coop, and Guy Sella. A genomic map of the effects of linked selection in drosophila. PLoS genetics, 12(8):e1006130, 2016.

[10] Guy Sella, Dmitri A Petrov, Molly Przeworski, and Peter Andolfatto. Pervasive natural selection in the drosophila genome? PLoS genetics, 5(6):e1000495, 2009.

[11] Asher D Cutter and Jae Young Choi. Natural selection shapes nucleotide polymorphism across the genome of the nematode caenorhabditis briggsae. Genome research, 20(8):1103–1111, 2010.

[12] Asher D Cutter and Bret A Payseur. Genomic signatures of selection at linked sites: unifying the disparity among species. Nature Reviews Genetics, 14(4):262–274, 2013.

[13] Josep M Comeron. Background selection as baseline for nucleotide variation across the drosophila genome. PLoS Genetics, 10(6):e1004434, 2014.

[14] Timothy M Beissinger, Li Wang, Kate Crosby, Arun Durvasula, Matthew B Hufford, and Jeffrey Ross-Ibarra. Recent demography drives changes in linked selection across the maize genome. Nature plants, 2(7):1–7, 2016.

[15] Raul Torres, Zachary A Szpiech, and Ryan D Hernandez. Human demographic history has amplified the effects of background selection across the genome. PLoS genetics, 14(6):e1007387, 2018.

[16] David Castellano, Adam Eyre-Walker, and Kasper Munch. Impact of mutation rate and selection at linked sites on dna variation across the genomes of humans and other homininae. Genome biology and evolution, 12(1):3550–3561, 2020.

[17] Gregory B Ewing and Jeffrey D Jensen. The consequences of not accounting for background selection in demographic inference. Molecular ecology, 25(1):135–141, 2016.

[18] Daniel R Schrider, Alexander G Shanku, and Andrew D Kern. Effects of linked selective sweeps on demographic inference and model selection. Genetics, 204(3):1207–1223, 2016.

[19] Parul Johri, Kellen Riall, Hannes Becher, Laurent Excoffier, Brian Charlesworth, and Jeffrey D Jensen. The impact of purifying and background selection on the inference of population history: problems and prospects. Molecular biology and evolution, 38(7):2986–3003, 2021.

[20] Simon Boitard, Armando Arredondo, Lounès Chikhi, and Olivier Mazet. Heterogeneity in effective size across the genome: effects on the inverse instantaneous coalescence rate (iicr) and implications for demographic inference under linked selection. Genetics, 220(3):iyac008, 2022.

[21] Laurent Excoffier, Isabelle Dupanloup, Emilia Huerta-Sánchez, Vitor C Sousa, and Matthieu Foll. Robust demographic inference from genomic and snp data. PLoS genetics, 9(10):e1003905, 2013.

[22] Richard R Hudson and Norman L Kaplan. Deleterious background selection with recombination. Genetics, 141(4):1605–1617, 1995.

[23] Magnus Nordborg, Brian Charlesworth, and Deborah Charlesworth. The effect of recombination on background selection. Genetics Research, 67(2):159–174, 1996.

[24] David Enard, Philipp W Messer, and Dmitri A Petrov. Genome-wide signals of positive selection in human evolution. Genome research, 24(6):885–895, 2014.

[25] 1000 Genomes Project Consortium et al. A global reference for human genetic variation. Nature, 526(7571):68, 2015.

[26] David A Murphy, Eyal Elyashiv, Guy Amster, and Guy Sella. Broad-scale variation in human genetic diversity levels is predicted by purifying selection on coding and non-coding elements. Elife, 12:e76065, 2022.

[27] Alan Hodgkinson and Adam Eyre-Walker. Variation in the mutation rate across mammalian genomes. Nature reviews genetics, 12(11):756–766, 2011.

[28] Hans Ellegren, Nick GC Smith, and Matthew T Webster. Mutation rate variation in the mammalian genome. Current opinion in genetics & development, 13(6):562–568, 2003.

[29] Benjamin C Haller and Philipp W Messer. Slim 3: forward genetic simulations beyond the wright–fisher model. Molecular biology and evolution, 36(3):632–637, 2019.

[30] Kay Prüfer, Fernando Racimo, Nick Patterson, Flora Jay, Sriram Sankararaman, Susanna Sawyer, Anja Heinze, Gabriel Renaud, Peter H Sudmant, Cesare De Filippo, et al. The complete genome sequence of a neanderthal from the altai mountains. Nature, 505(7481):43–49, 2014.

[31] Marta Byrska-Bishop, Uday S Evani, Xuefang Zhao, Anna O Basile, Haley J Abel, Allison A Regier, André Corvelo, Wayne E Clarke, Rajeeva Musunuri, Kshithija Nagulapalli, et al. High-coverage wholegenome sequencing of the expanded 1000 genomes project cohort including 602 trios. Cell, 185(18):3426–3440, 2022.

[32] Débora YC Brandt, Xinzhu Wei, Yun Deng, Andrew H Vaughn, and Rasmus Nielsen. Evaluation of methods for estimating coalescence times using ancestral recombination graphs. Genetics, 221(1):iyac044, 2022.

[33] Regev Schweiger and Richard Durbin. Ultra-fast genome-wide inference of pairwise coalescence times. bioRxiv, pages 2023–01, 2023.

[34] Seong-Ho Kim, Navin Elango, Charles Warden, Eric Vigoda, and Soojin V Yi. Heterogeneous genomic molecular clocks in primates. PLoS genetics, 2(10):e163, 2006.

[35] Jedidiah Carlson, Adam E Locke, Matthew Flickinger, Matthew Zawistowski, Shawn Levy, Richard M Myers, Michael Boehnke, Hyun Min Kang, Laura J Scott, Jun Z Li, et al. Extremely rare variants reveal patterns of germline mutation rate heterogeneity in humans. Nature communications, 9(1):3753, 2018.

[36] Varun Aggarwala and Benjamin F Voight. An expanded sequence context model broadly explains variability in polymorphism levels across the human genome. Nature genetics, 48(4):349–355, 2016.

[37] Christopher J Adams, Mitchell Conery, Benjamin J Auerbach, Shane T Jensen, Iain Mathieson, and Benjamin F Voight. Regularized sequence-context mutational trees capture variation in mutation rates across the human genome. PLoS Genetics, 19(7):e1010807, 2023.

[38] Leo Speidel, Marie Forest, Sinan Shi, and Simon R Myers. A method for genome-wide genealogy estimation for thousands of samples. Nature genetics, 51(9):1321–1329, 2019.

[39] Matthew D Rasmussen, Melissa J Hubisz, Ilan Gronau, and Adam Siepel. Genome-wide inference of ancestral recombination graphs. PLoS genetics, 10(5):e1004342, 2014.

[40] Olivier Mazet, Willy Rodríguez, and Lounès Chikhi. Demographic inference using genetic data from a single individual: Separating population size variation from population structure. Theoretical population biology, 104:46–58, 2015.

[41] Olivier Mazet, Willy Rodríguez, Simona Grusea, Simon Boitard, and Lounès Chikhi. On the importance of being structured: instantaneous coalescence rates and human evolution—lessons for ancestral population size inference? Heredity, 116(4):362–371, 2016.

[42] John E Pool and Rasmus Nielsen. Population size changes reshape genomic patterns of diversity. Evolution, 61(12):3001–3006, 2007.

[43] Guy Amster, David A Murphy, William R Milligan, and Guy Sella. Changes in life history and population size can explain the relative neutral diversity levels on x and autosomes in extant human populations. Proceedings of the National Academy of Sciences, 117(33):20063–20069, 2020.

[44] Guy Amster and Guy Sella.Life history effects on neutral diversity levels of autosomes and sex chromosomes. Genetics, 215(4):1133–1142, 2020.

[45] Guy Amster and Guy Sella. Life history effects on the molecular clock of autosomes and sex chromosomes. Proceedings of the National Academy of Sciences, 113(6):1588–1593, 2016.

[46] Jerome Kelleher, Yan Wong, Anthony W Wohns, Chaimaa Fadil, Patrick K Albers, and Gil McVean. Inferring whole-genome histories in large population datasets. Nature genetics, 51(9):1330–1338, 2019.

[47] Anastasia Ignatieva, Rune B Lyngsø, Paul A Jenkins, and Jotun Hein. Kwarg: Parsimonious recon-struction of ancestral recombination graphs with recurrent mutation. Bioinformatics, 37(19):3277–3284, 2021.

[48] Anthony Wilder Wohns, Yan Wong, Ben Jeffery, Ali Akbari, Swapan Mallick, Ron Pinhasi, Nick Patterson, David Reich, Jerome Kelleher, and Gil McVean. A unified genealogy of modern and ancient genomes. Science, 375(6583):eabi8264, 2022.

[49] Brian C Zhang, Arjun Biddanda, Árni Freyr Gunnarsson, Fergus Cooper, and Pier Francesco Palamara. Biobank-scale inference of ancestral recombination graphs enables genealogical analysis of complex traits. Nature Genetics, pages 1–9, 2023.

[50] Lauren E Nicolaisen and Michael M Desai. Distortions in genealogies due to purifying selection and recombination. Genetics, 195(1):221–230, 2013.

[51] Yiyuan Fang, Shuyi Deng, and Cai Li. A generalizable deep learning framework for inferring fine-scale germline mutation rate maps. Nature Machine Intelligence, 4(12):1209–1223, 2022.

[52] Thomas CA Smith, Peter F Arndt, and Adam Eyre-Walker. Large scale variation in the rate of germ-line de novo mutation, base composition, divergence and diversity in humans. PLoS genetics, 14(3):e1007254, 2018.

[53] Ivana Cvijović, Benjamin H Good, and Michael M Desai. The effect of strong purifying selection on genetic diversity. Genetics, 209(4):1235–1278, 2018.

[54] Vince Buffalo and Andrew D Kern. A quantitative genetic model of background selection in humans. bioRxiv, pages 2023–09, 2023.

[55] Søren Besenbacher, Christina Hvilsom, Tomas Marques-Bonet, Thomas Mailund, and Mikkel Heide Schierup. Direct estimation of mutations in great apes reconciles phylogenetic dating. Nature ecology & evolution, 3(2):286–292, 2019.

[56] Angela S Hinrichs, Donna Karolchik, Robert Baertsch, Galt P Barber, Gill Bejerano, Hiram Clawson, Mark Diekhans, Terrence S Furey, Rachel A Harte, Fan Hsu, et al. The ucsc genome browser database: update 2006. Nucleic acids research, 34(suppl 1):D590–D598, 2006.

[57] Aylwyn Scally and Richard Durbin. Revising the human mutation rate: implications for understanding human evolution. Nature Reviews Genetics, 13(10):745–753, 2012.

[58] Whole-Genome Sequencing. Analysis of genetic inheritance in a family quartet by. Nature, 432:695, 2004.

[59] 1000 Genomes Project Consortium et al. A map of human genome variation from population scale sequencing. Nature, 467(7319):1061, 2010.

[60] Philip Awadalla, Julie Gauthier, Rachel A Myers, Ferran Casals, Fadi F Hamdan, Alexander R Griffing, Mélanie Côté, Edouard Henrion, Dan Spiegelman, Julien Tarabeux, et al. Direct measure of the de novo mutation rate in autism and schizophrenia cohorts. The American Journal of Human Genetics, 87(3):316–324, 2010.

[61] Jack N Fenner. Cross-cultural estimation of the human generation interval for use in genetics-based population divergence studies. American Journal of Physical Anthropology: The Official Publication of the American Association of Physical Anthropologists, 128(2):415–423, 2005.

[62] Manjusha Chintalapati and Priya Moorjani. Evolution of the mutation rate across primates. Current opinion in genetics & development, 62:58–64, 2020.

[63] Lawrence Rabiner and Biinghwang Juang. An introduction to hidden markov models. ieee assp magazine, 3(1):4–16, 1986.

[64] Richard Durbin, Sean R Eddy, Anders Krogh, and Graeme Mitchison. Biological sequence analysis: probabilistic models of proteins and nucleic acids. Cambridge university press, 1998.

[65] Christopher M Bishop and Nasser M Nasrabadi. Pattern recognition and machine learning, volume 4. Springer, 2006.

[66] Bernard Y Kim, Christian D Huber, and Kirk E Lohmueller. Inference of the distribution of selection coefficients for new nonsynonymous mutations using large samples. Genetics, 206(1):345–361, 2017.

[67] Aylwyn Scally. The mutation rate in human evolution and demographic inference. Current opinion in genetics & development, 41:36–43, 2016.

[68] Nick Patterson, Daniel J Richter, Sante Gnerre, Eric S Lander, and David Reich. Genetic evidence for complex speciation of humans and chimpanzees. Nature, 441(7097):1103–1108, 2006.

